# Extracellular stiffness regulates cell fate determination and drives the emergence of evolutionary novelty in teleost heart

**DOI:** 10.1101/2025.05.13.653630

**Authors:** Sho Matsuki, Yusei Inoue, Ryuta Watanabe, Toshiyuki Mitsui, Yuuta Moriyama

## Abstract

The diversification of life is driven by the acquisition of new phenotypic traits, known as evolutionary novelties. While some genetic mechanisms underlying such traits have been identified, the role of physical properties of cellular and extracellular components remains largely unexplored. Here, we show that the evolution and development of the bulbus arteriosus (BA)–a teleost-specific, smooth muscle-rich heart outflow tract–is regulated by the mechanical properties of its extracellular environment. Specifically, we demonstrate that the teleost-specific extracellular matrix gene *elastin b* confers the uniquely low stiffness of the BA. In contrast to the BA, the homologous organ in non-teleost actinopterygians, the conus arteriosus (CA), and the ventricle are predominantly composed of cardiomyocytes, and exhibit higher extracellular stiffness. Loss of *elastin b* function by knockdown or knockout leads to ectopic cardiomyocyte formation in the BA, accompanied by increased extracellular stiffness comparable to that of the ventricle. Furthermore, artificial stiffening of the BA extracellular environment is sufficient to induce ectopic cardiomyocyte differentiation. Taken together, these findings demonstrate that extracellular stiffness governs cell fate determination and highlight its role in the emergence of evolutionary novelties in the teleost heart.

**Significant Statement:** The acquisition of new phenotypic traits is a key driver for a diversification of life. While genetic underpinnings of such traits have been well studied, the contribution of physical properties remains elusive. Here, we show that extracellular stiffness plays a key role in the evolution and development of the bulbus arteriosus (BA), a smooth muscle-based cardiac outflow tract unique to teleost hearts. We demonstrate that *elastin b*, a teleost-specific gene, confers characteristically low stiffness to the BA, which is required for proper cell fate determination. Notably, such low stiffness is absent in non-teleost fishes and in teleosts with impaired *elastin b* function, and artificial stiffening alters cell fate. Our findings highlight the importance of physical properties in driving evolutionary novelties.

## Introduction

The diversification of life is driven by the *de novo* evolution of complex phenotypic traits known as evolutionary novelties. While genetic mechanisms underlying the morphogenesis of evolutionary novelties during embryonic development have been extensively studied (1), the role of physical properties of both cellular or non-cellular components in these processes remains largely unexplored.

A component of the teleost heart, the bulbus arteriosus (BA), represent one of the most important evolutionary novelties in the teleost lineage (2). The BA is a specialized heart outflow tract (OFT) that plays a critical role in the Windkessel function–absorbing the pulsatile pressure and flow generated by the ventricle through elastic expansion and recoil, thereby smoothing these parameters to ensure stable perfusion through the downstream arterial system (3–11). The BA also exhibits unique anatomical features in its cellular composition: it is primarily composed of smooth muscle, whereas the OFTs of other vertebrates are mainly composed of cardiac muscle with contractile properties. This anatomical uniqueness is thought to be essential for the functional specialization of the BA.

We previously demonstrated that *elastin b* (*elnb*), a teleost-specific extracellular matrix (ECM) gene, plays a pivotal role in both the evolution and development of the teleost BA (12). Loss-of-function of *elnb* resulted in ectopic cardiomyocytes within the BA without changes of cardiac precursors’ migration pattern. Furthermore, knockdown of *yap*—a mechanotransducer that senses and translates extracellular physical properties into intracellular biochemical signaling—also led to ectopic cardiomyocyte formation in the BA without altering *elnb* expression. These findings suggest that Elastin b (Elnb) possesses unique physical properties such as stiffness and endows specific extracellular environment for the BA, and cells in the BA sense such physical information via mechanotransduction by *yap* and specify their own cell fate as smooth muscle. However, the physical properties of extracellular environment in the BA and other cardiac components have not been fully characterized yet.

The ECM is a non-cellular component that fills the extracellular space within tissues and organs. While ECM has long been considered as a structural scaffold or substrate for cell migration, recent studies have revealed that the ECM’s physical properties, such as stiffness and viscoelasticity, also plays an important role in various biological processes, including cell growth, proliferation, fate determination, migration, immunity, malignant transformation and apoptosis (13, 14). In particular, the role of the ECM in cell fate determination has received considerable attention in recent years in the context of regenerative medicine and is actively investigated using *in vitro* systems such as organoid models (15–18). However, the contribution of ECM physical properties to cell fate determination *in vivo*–during the processes such as embryonic development (ontogeny) and evolutionary diversification (phylogeny)–remains largely unknown.

Atomic force microscopy (AFM) is one of the most effective and widely used instruments for measuring physical properties such as stiffness and elasticity in various materials. In biological systems, AFM has been adapted to measure relatively soft, elastic materials through modifications such as attaching micrometer-sized beads to the tip of cantilever (19) and applying appropriate mechanical models like the Hertz model to extract parameters (20). AFM-based approaches have revealed a broad range of mechanical and physical properties in living systems, both *in vitro* and *in vivo*, and have become central to the field of mechanobiology, which involves biology, medicine, biophysics and bioengineering.

Here, we describe the mechanical properties of extracellular environment in vertebrate heart components, with a particular focus on the BA in teleosts, the conus arteriosus (CA)—the homologous outflow tract in non-teleost fishes—and the ventricle. In teleosts (zebrafish, *Danio rerio*), we found that the extracellular environment of the BA exhibits significantly lower stiffness compared to that of the ventricle. In contrast, in non-teleost fish (*Polypterus*, *Polypterus senegalus*), the CA and the ventricle showed similar stiffness levels. Furthermore, in *elnb* knockdown (KD) and knockout (KO) zebrafish, both of which develop ectopic cardiomyocytes in the BA, we observed that the stiffness of the extracellular environment in the regions where ectopic cardiomyocytes formed was increased and comparable to that of the ventricle. A comparison of OFT stiffness at 2 days post fertilization (dpf), prior to *elnb* expression, and BA stiffness at 4 dpf, after *elnb* expression begins, revealed that the characteristic lower stiffness emerged specifically in the BA at 4 dpf, suggesting that *elnb* contributes to the mechanical properties of the BA extracellular environment. Finally, we artificially increased the stiffness of the BA extracellular environment by applying local mechanical pressure using a glass capillary, and this manipulation led to the formation of ectopic cardiomyocytes in the BA, phenocopying the *elnb* KD/KO condition. These findings suggest that the extracellular stiffness of the BA plays a crucial role in cell fate determination and is essential for the proper development of the BA during teleost heart formation. Our study highlights the functional significance of physical properties–specifically extracellular stiffness–in the evolution and development of the teleost heart.

## Results

### Extracellular environment in the BA exhibits uniquely low stiffness compared with that of the ventricle in the teleost heart

We hypothesized that the difference in cell composition between the BA and other heart components in teleosts–smooth muscle-dominant in the BA and cardiomyocyte-dominant elsewhere (Fig. 1*A*)—may be determined by distinct ECM stiffness, given that a teleost-specific ECM gene *elnb* is expressed in the BA and its loss results in ectopic cardiomyocytes within this structure (12). To test this hypothesis, we characterized the extracellular stiffness of teleost heart components using atomic force microscopy (AFM) in combination with decellularization techniques.

**Fig. 1.**
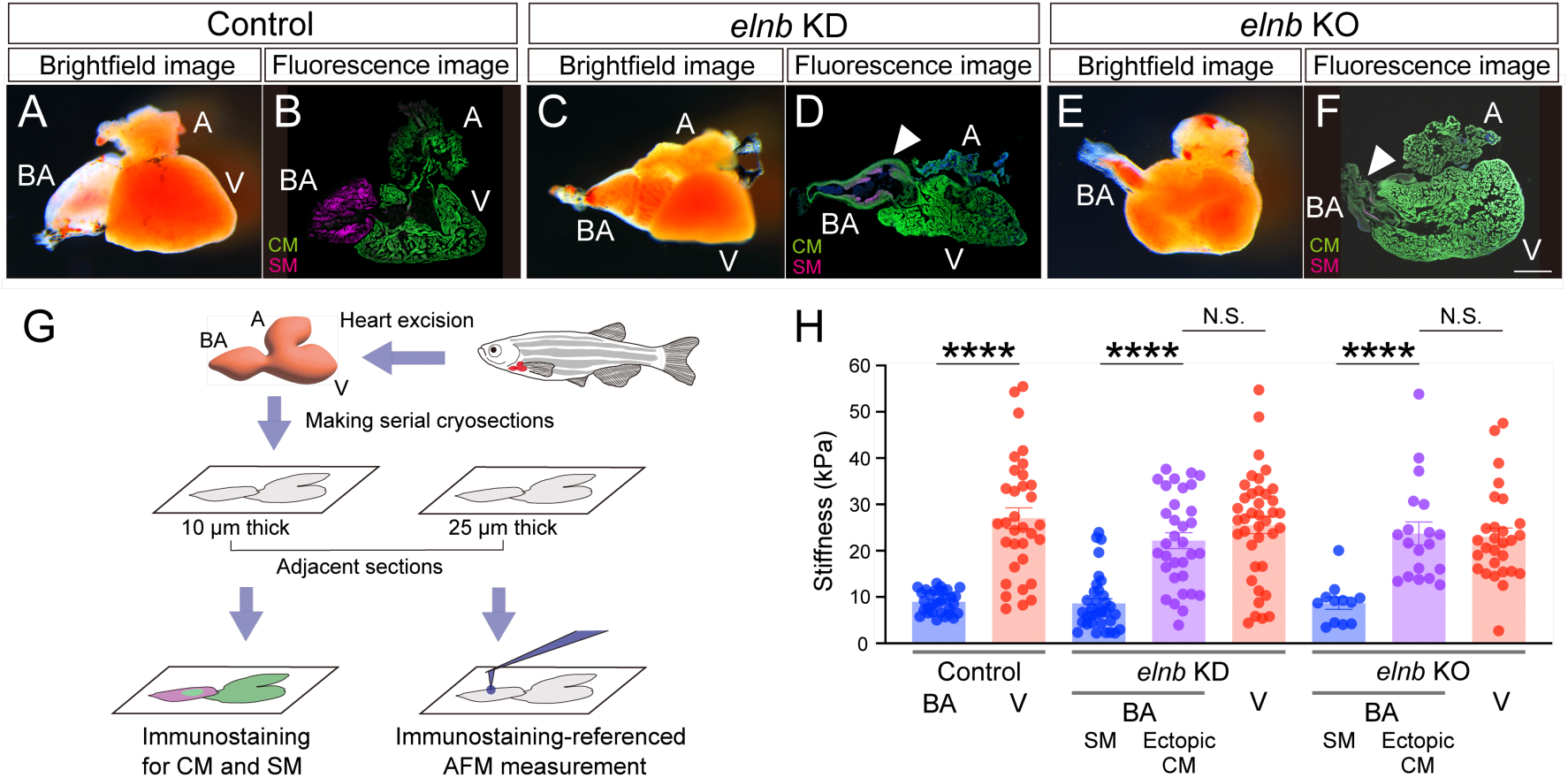
Heat morphology, cellular composition, and extracellular stiffness in adult control, *elnb* knockdown, and *elnb* knockout zebrafish hearts. (*A-F*) Brightfield images of heart morphology and fluorescence images of immunostained sections of control (*A* and *B*), *elnb* knockdown (KD; *C* and *D*) and *elnb* knockout (KO; *E* and *F*) zebrafish. Green, cardiomyocytes; Magenta, smooth muscle. Scale bar, 200 µm. Arrowheads indicate ectopic cardiomyocytes formed in the BA. (*G*) Schematic representation of AFM-based extracellular stiffness measurement, guided by immunostaining of adjacent sections. (*H*) Quantification of extracellular stiffness in the BA and ventricle (V) of control hearts (BA: n=31; V: n=33), and in *elnb* KD and KO hearts. In *elnb* KD hearts, measurements were performed in smooth muscle-forming regions (BA-SM; n=34) and ectopic cardiomyocyte-forming regions (BA-CM; n=34) in the BA, and the V (n=39). In *elnb* KO hearts, measurements were performed in BA-SM (n=12) and BA-CM (n=19), and the V (n=28). Statistical significances were assessed using the Mann–Whitney test for control, and the Kruskal–Wallis test for *elnb* KD and KO groups. N.S., *P*>0.05; ****, *P*<0.0001. Data represent means ±s.e.m. A, atrium; AFM, atomic force microscopy; BA, bulbus arteriosus; CM, cardiomyocyte; SM, smooth muscle; V, ventricle.

Decellularization allows for the removal of all cellular components, leaving only the ECM or other non-cellular structures for mechanical measurement (Fig. S1*A*). In recent years, various decellularization methods have been developed and are now widely used in tissue and ECM studies (21). Among these, we employed a sodium deoxycholate (SDC)-based protocol, which offers superior preservation of ECM integrity following treatment (22).

We first characterized the extracellular stiffness of the BA and ventricle in the adult zebrafish (teleost) heart. Hearts containing both the BA and ventricle were surgically dissected and sectioned into alternating adjacent slices of 10 µm and 25 µm thickness, which were used for immunostaining and AFM measurements, respectively.

Immunostaining was performed to identify regions within the BA that contained either smooth muscle or cardiomyocytes. Using the adjacent sections, we then referred to the staining results to guide AFM-based measurements of the Young’s modulus in the corresponding smooth muscle and cardiomyocyte regions. AFM samples were subjected to decellularization prior to AFM analysis. As a result, we found that the effective Young’s modulus (hereafter referred to as “stiffness”) of the BA and ventricle were 8.98 (± 0.43) and 27.04 (± 2.24) kPa, respectively, with a significant difference between the two regions (*p* < 0.0001; Fig. 1*D*).

Next, we assessed extracellular stiffness in *elnb* KD adult zebrafish (morphants), which have previously been shown to develop ectopic cardiomyocytes in the BA (12)(Fig. 1*B*). We measured the extracellular stiffness in the *elnb* morphants’ BA containing either smooth muscle or ectopic cardiomyocytes, and found values of 8.63 (± 1.04) and 23.73 (± 2.28) kPa, respectively, again showing a significant difference between these (*p* < 0.0001; Fig. 1*D*). Notably, the stiffness in the region of ectopic cardiomyocytes in *elnb* morphants was comparable to that of the ventricle (25.56 ± 1.85 kPa), with no significant difference between the two (*p* > 0.05; Fig. 1*D*).

We then analyzed *elnb* KO zebrafish generated by CRISPR/Cas9 system. This *elnb* mutant carries a premature stop codon that results in a truncated 81-amino acid Elnb protein, compared to the full length 2,041-amino acid protein in wild-type fish (Fig. S1*B*). Homozygous *elnb* mutants exhibited a hypoplastic BA and developed ectopic cardiomyocytes in the BA (Fig. 1*C*). The stiffness in BA regions containing smooth muscle and ectopic cardiomyocytes were 8.64 (± 1.31) and 23.73 (± 2.47) kPa, respectively (*p* < 0.0001; Fig. 1*D*), while the ventricle showed a stiffness of 23.03 (± 1.87) kPa. As in the morphants, the stiffness in the region with ectopic cardiomyocytes did not differ significantly from that of the ventricle (*p* > 0.05; Fig. 1*D*). Taken together, these results demonstrate a strong correlation between extracellular stiffness and cell type determination in zebrafish heart components.

### Extracellular environment in the ventricle and CA of adult *Polypterus* exhibits similar stiffness

Our results from adult zebrafish hearts suggest a correlation between cell type (smooth muscle vs. cardiomyocytes) and extracellular stiffness in the BA, and highlight a role for *elnb* in mediating this relationship. To further investigate this, we next characterized the extracellular stiffness of the CA and ventricle in adult *Polypterus*, a representative non-teleost actinopterygian. The CA in *Polypterus* is primarily composed of cardiomyocytes, similar to the ventricle and other cardiac regions (2)(Fig. 2*A*). We found that extracellular stiffness of the CA and ventricle in *Polypterus* were 26.22 (± 1.25) and 26.33 (± 1.75) kPa, respectively. Notably, there was no significant difference between these two regions (*p* > 0.05; Fig. 2*B*), in contrast to the marked stiffness difference between the BA and ventricle observed in teleosts. These findings suggest that in the absence of *elnb*, the stiffness of the OFT remains comparable to that of the ventricle in non-teleost actinopterygians—similar to what is observed in *elnb* KD/KO zebrafish, where ectopic cardiomyocytes are formed in the BA. This supports the idea that the teleost-specific ECM component Elnb imparts distinct mechanical properties to the BA extracellular environment, facilitating its unique smooth muscle-rich identity.

**Fig. 2.**
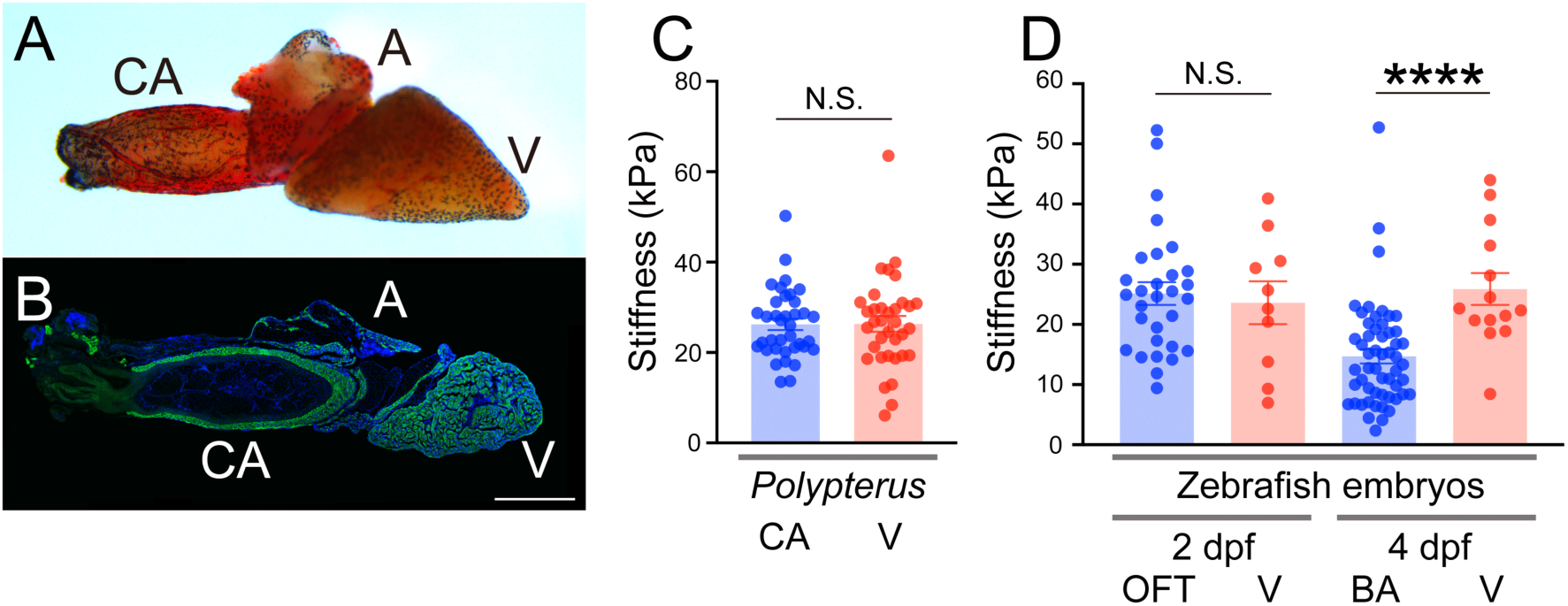
Heart morphology and cellular composition in adult *Polypterus*, and extracellular stiffness in adult *Polypterus* and developing zebrafish hearts. (*A* and *B*) Brightfield image of heart morphology (*A*) and fluorescence image of an immunostained section (*B*) of adult *Polypterus* heart. Scale bar, 600 µm. (*C*) Quantification of extracellular stiffness in the CA and V of adult *Polypterus* (CA: n=35; V: n=37). Statistical significance was assessed using Mann–Whitney test. N.S., *P*>0.05. (*D*) Quantification of extracellular stiffness in the OFT and V in 2 dpf zebrafish embryos (OFT: n=30; V: n=10), and in the BA and V in 4 dpf embryos (BA: n=54; V: n=14). Statistical significances were assessed using Mann–Whitney test. N.S., *P*>0.05; ****, *P*<0.0001. Data represent means ±s.e.m. A, atrium; BA, bulbus arteriosus; CA, conus arteriosus; OFT, outflow tract; V, ventricle.

### Unique extracellular stiffness in the BA emerges after the initiation of *elnb* expression

Our characterization of extracellular stiffness in adult zebrafish and *Polypterus* hearts suggests that *elnb* contributes to the distinct stiffness observed in the BA. To test this, we next examined extracellular stiffness in the OFT/BA and ventricle of developing zebrafish embryos. Previous studies have shown that *elnb* expression begins at 3 days post-fertilization (dpf), specifically in the OFT (future BA region), and that smooth muscle cells appear in the BA from 4 dpf onward (12). Therefore, we measured extracellular stiffness in the OFT/BA and ventricle at 2 and 4 dpf, corresponding to stages before and after *elnb* expression in the OFT/BA. At 2 dpf, the extracellular stiffness of the OFT and ventricle was 25.13 (± 1.89) and 23.60 (± 3.56) kPa, respectively, with no significant difference between the two (*p* > 0.05; Fig. 2*C*). In contrast, at 4 dpf, the extracellular stiffness of the BA and ventricle were 14.69 (± 1.18) and 25.88 (± 2.74) kPa, respectively, with the BA showing a marked reduction in extracellular stiffness and a significant difference from the ventricle, similar to that observed in adult zebrafish hearts (*p* < 0.0001; Fig. 2*C*). These findings indicate that the differential stiffness between the OFT/BA and the ventricle arises after the onset of *elnb* expression and support the idea that *elnb* contributes to establishing the unique mechanical properties of the BA extracellular environment.

### Artificial stiffening of the extracellular environment in the BA induces ectopic cardiomyocyte formation

Lastly, to further investigate the relationship between extracellular stiffness and cell fate determination, we artificially increased the stiffness of the OFT/BA region and examined its effect on cardiac cell fate. A previous study in *Xenopus* embryos employed a local indentation method to increase substrate stiffness and revealed underlying mechanisms of collective durotaxis (23). Following a similar approach, we applied localized mechanical pressure to the OFT/BA region using a glass capillary.

Indentation was initiated at 60 hours post-fertilization (hpf) and maintained until 96 hpf. Embryos were then fixed and analyzed for the presence of ectopic cardiomyocytes in the BA (Fig. 3*A*). As a result, we observed ectopic cardiomyocyte formation in the BA in 71.4% of indented embryos (n = 5/7), whereas no control embryos—embedded in low-melting-point agarose without indentation—exhibited such ectopic cardiomyocytes in the BA (n = 0/8; Fig. 3*B* and *C*). Notably, *elnb* expression in the BA of indented embryos remained comparable to that of controls (Fig. 3*D* and *E*). These findings suggest that increased extracellular stiffness in the OFT/BA region is sufficient to induce cardiomyocyte differentiation, supporting a role for mechanical cues in regulating cell fate during BA morphogenesis in zebrafish development.

**Fig. 3.**
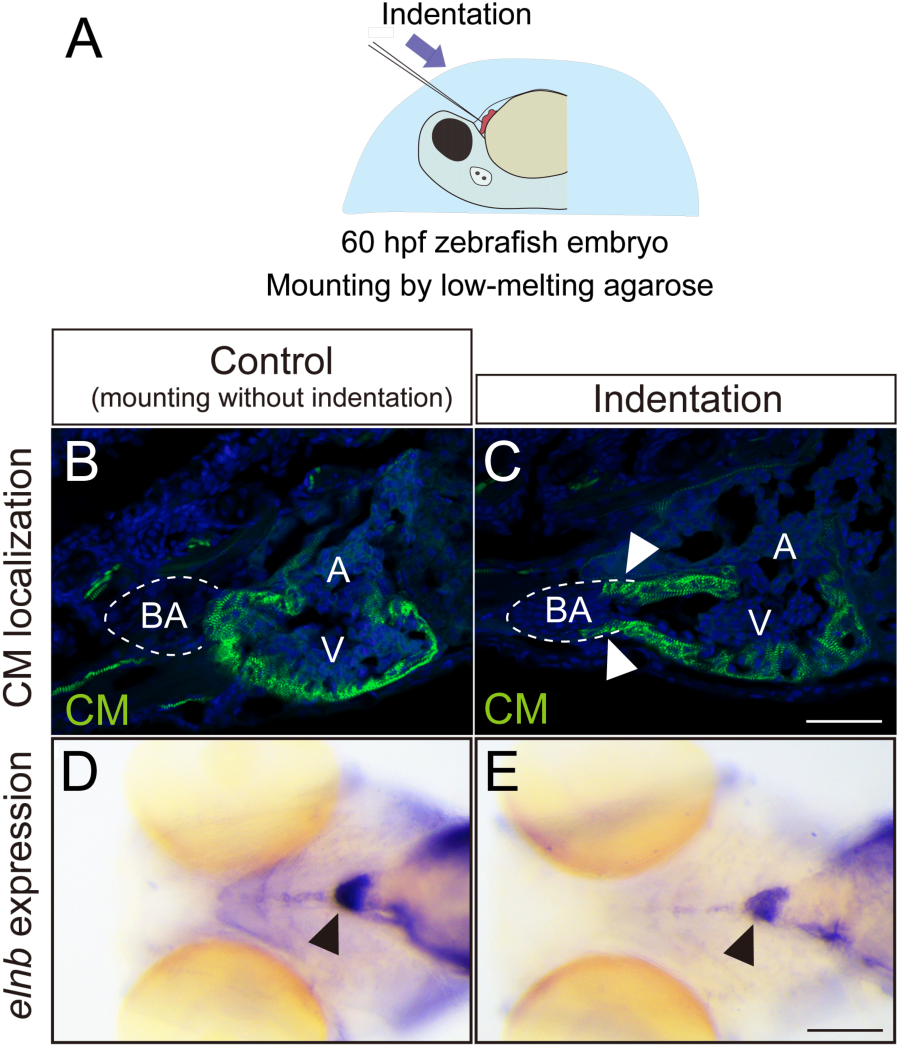
Artificial stiffening of the extracellular environment induces ectopic cardiomyocyte formation in the BA of developing zebrafish. (A) Schematic diagram of the indentation experiment. Local indentation was performed using a glass needle targeting the developing OFT in 60 hpf zebrafish embryos. (*B* and *C*) Fluorescence images of immunostained sections of control (mounting without indentation; *B*, n = 0/8) and indented (*C*, n = 5/7) 96 hpf zebrafish hearts. Green, cardiomyocytes. Scale bar, 50 µm. Arrowheads indicate ectopic cardiomyocytes formed in the BA. (*D* and *E*) Expression patterns of *elnb* in control (*D*) and indented (*E*) 4 dpf zebrafish embryos. Scale bar, 200 µm. A, atrium; BA, bulbus arteriosus; CM, cardiomyocytes; V, ventricle.

## Discussion

Here, we characterized extracellular stiffness in the ventricle and OFT—specifically, the BA in teleosts and CA in non-teleost fishes—and identified a correlation between the expression of the teleost-specific ECM-coding gene *elnb* and reduced extracellular stiffness. In teleost, *elnb* is specifically expressed in the BA, where the extracellular environment exhibits a uniquely low stiffness compared to the ventricle or to the CA in non-teleost fishes (Fig. 1*H*; Fig. 2*C*; 12). During zebrafish embryogenesis, this characteristic reduction in stiffness becomes detectable at 4 dpf, following the onset of *elnb* expression at 3 dpf, and is not observed at 2 dpf (Fig. 2*D*; 12). Furthermore, in *elnb* KD/KO embryos— where ectopic cardiomyocytes form within the BA—the stiffness of the extracellular environment in regions containing these ectopic cells was increased to levels comparable to that of the ventricle (Fig. 1*H*). These findings suggest that Elnb intrinsically confers a uniquely low stiffness to the extracellular environment and promotes the differentiation of cardiac precursor cells into smooth muscle in the BA. Consistent with this idea, recent studies have increasingly shown that ECM stiffness can influence stem cell fate determination (13, 24, 25). However, most of this evidence comes from *in vitro* systems, and whether ECM or extracellular stiffness directs cell fate decisions during embryonic development—or contributes to the emergence of evolutionary novelties—remains largely unexplored. Our findings provide new insight into this question: the teleost-specific ECM component (Elnb) may guide the differentiation of cells in the OFT/BA toward a smooth muscle fate, contributing not only to BA morphogenesis (ontogeny) but also to the evolutionary acquisition of the BA in teleosts (phylogeny).

Zebrafish possess a remarkable ability to regenerate their heart after injury, and the role of the ECM in this process has been extensively studied. Distinct ECM components have been identified across different regions of the zebrafish heart, and during ventricular regeneration, the ECM undergoes dynamic remodeling. A significant reduction in ECM stiffness has been reported at 7 days post-amputation, not only at the site of injury but also in remote cardiac regions (26). Moreover, a previous study demonstrated that low ECM stiffness serves as a permissive cue for heart regeneration in neonatal mice (27). In support of this, genetic ablation of *collagen 1 alpha 2* (*col1a2*)-expressing cells in injured hearts impaired cardiomyocyte proliferation during the regenerative process (28). These findings collectively highlight the importance of ECM mechanical properties in cardiac regeneration. Interestingly, the BA has also been shown to play an essential role in the regeneration of the adjacent ventricular epicardium. Grafting the BA into an injured ventricle was found to initiate and redirect epicardial regeneration through Hedgehog signaling (29). Based on these observations, it would be of great interest to investigate whether the BA-specific ECM component Elnb contributes to cardiac regeneration in teleosts.

The BA is considered one of the most significant evolutionary novelties in teleosts, functioning as a Windkessel organ that undergoes elastic expansion and recoil to absorb and dampen the pulsatile blood flow ejected from the ventricle (3–11). Numerous studies have shown that the BA in various teleost species is highly distensible and resilient, and that these mechanical properties are essential for its Windkessel function (3, 4, 6, 8, 10). Intriguingly, we previously reported that *elnb* KD in zebrafish embryos resulted in reduced deformation amplitude and shortened duration of BA contraction (12). Given the correlation between *elnb* expression and extracellular stiffness identified in the present study, it is plausible that Elnb contributes to the soft (i.e., low-stiffness) extracellular environment of the BA, and that this unique mechanical property is critical for enabling its Windkessel function.

In non-teleost vertebrates (e.g., mammals, birds, and reptiles), the *Elastin* gene exists as a single-copy gene without paralogues or sister genes in the genome, and encodes Tropoelastin, the monomeric precursor of Elastin. Secreted Tropoelastin forms microaggregates and, through interactions with components such as LTBP-4 and Fbln5, assembles into elastic fibers that provide stretchability, resilience, and cell-matrix interactions in various elastic tissues, including the skin, lungs, cardiovascular system, cartilage, and tendons (30, 31). While the molecular mechanisms of elastic fiber assembly have been extensively studied in mammals, how Elnb (Tropoelastin b) assembles and functions in teleosts remains largely unknown. Understanding how Elnb is assembled and what cofactors are required for its function represents an important avenue for future research. In this context, *ltbp3* emerges as a promising candidate, as it is specifically expressed in the developing BA and has been identified as a marker of the second heart field. Notably, *ltbp3* KD in zebrafish results in ectopic cardiomyocyte formation in the BA, phenocopying the *elnb* KD phenotype (12, 32). Elucidating the molecular mechanisms of Elnb assembly, as well as the evolutionary origins of this teleost-specific gene and its integration into pre-existing molecular networks, represent intriguing and open questions for future investigation.

## Materials and methods

### Zebrafish and *Polypterus* Husbandry

Zebrafish (*Danio rerio*) maintenance, microinjection, and staging were performed as previous described (33). The RW strain was used as the wild-type control. Embryos were obtained through natural spawning, maintained in E3 medium at 28.5 °C - 31 °C, and staged based on established morphological criteria (33). *Polypterus* (*Polypterus senegalus*) maintenance was conducted as describe previously (34). Adult *Polypterus* were obtained from Meito-suien (Aichi, Japan).

### Morpholino Injection

Zebrafish embryos were injected at one-cell stage using glass capillary needles (30-0020, Harvard Apparatus), which were pulled using a needle puller (P-97 IVF, Sutter Instrument) and mounted on a microinjection system (FemtoJet 4i, Eppendorf). Antisense morpholino oligonucleotide (MO) was designed to block translation and purchased from Gene Tools, LLC (Philomath, OR, USA). The sequence of *elnb* MO used in this study is as follows: (5’-CCGGGCCATCCTGCTCTGTAATAAC-3’). Each embryo was injected with 1 ng of MO.

### CRISPR/Cas9 Mutant Generation

The *elnb* KO mutant line was generated using CRISPR/Cas9 as previously described (35). The target site for *elnb* (ENSDARG00000069017) was determined using the CHOPCHOP website (http://chopchop.cbu.uib.no) (36, 37). To generate gRNA, the following oligonucleotide primer set was designed and used: *elnb* (5’-<u>TAGG</u>ggCATTGGAACAGGTCCTGG-3’, 5’-<u>AAAC</u>CCAGGACCTGTTCCAATGcc-3’). The target site was designed in the second exon of *elnb* gene. In the above gRNA oligonucleotide sequences, underlined regions correspond to sequences added for subcloning into the BsaI-digested pDR274 plasmid, and lowercase letters indicate bases added to enhance *in vitro* gRNA transcription efficiency. To anneal the gRNA oligonucleotides and generate linker DNA, 9 µl of each 100 µM oligonucleotide and 2 µl of 10x M buffer (TAKARA) were mixed and incubated at 72 °C for 10 minutes, followed by incubation at room temperature for 20 minutes. Then, 180 µl of TE buffer was added, and the mixture was placed on ice. The annealed linker DNAs were cloned into BsaI-digested pDR274 plasmid, and the resulting plasmid was digested with HindIII. *In vitro* transcription of gRNA was performed using MEGAshortscript T7 Transcription kit (Ambion). For microinjection, 1 - 2 nl of a mixture of 0.2 µl of 400 ng/µl gRNA, 2 µl of 300 ng/µl Cas9 mRNA, 0.2 µl of 400 ng/µl tyrosinase gRNA, and 0.2 µl of 1M KCl was injected into one-cell stage zebrafish embryos. Tyrosinase gRNA was used for G0 screening (38). Cas9 mRNA was synthesized using mMESSAGE mMACHINE SP6 Transcription Kit (Ambion). For screening of mutant fish with germline transmission, heteroduplex mobility assay (HMA) was used (39). Primer sequences used for HMA are as follows: (5’-AATTTAGGTTATGTTGGGCCTG-3’, 5’-GCAAAGTAAATGCATCCCTACC-3’).

For the identification of mutated sequences, the amplified PCR products were cloned into the Zero Blunt TOPO PCR vector and sequenced (Eurofins Genomics). Identified mutant *elnb* fish were outcrossed with wild-type fish to generate heterozygous lines. Genotyping of *elnb* mutant embryos was performed by PCR using same primer set as used for HMA.

### Cryosectioning

Embryos or extracted adult hearts were transferred into phosphate-buffered saline (PBS) containing increasing concentrations of sucrose (5%, 15%, 30% [w/v] sucrose in PBS), with incubation at 4 °C overnight for each step. Samples were then incubated in a 1:1 mixture of 30% sucrose/PBS and optimal cutting temperature compound (OCT; Tissue-Tek) for 2 hours at 4 °C. Subsequently, samples were transferred into disposable embedding molds (Cryomold; Tissue-Tek) filled with OCT and frozen at-80°C. Frozen samples embedded in OCT were sectioned into alternating adjacent slices of 10 µm and 25 µm thickness using a Leica CM 3050 S cryostat; the former were used for immunostaining, and the latter for immunostaining-referenced AFM measurements.

### Atomic Force Microscopy

For the measurement of the extracellular stiffness in zebrafish (embryonic and adult) and *Polypterus* (adult) hearts, samples were prepared by cryosectioning at 25 µm thickness in OCT compound and mounted on MAS-coated glass slides (MATSUNAMI). Prior to measurement, samples were washed with PBS and decellularized to remove cellular components with preserving ECM components. AFM measurements performed immediately after decellularization using a scanning probe microscope (SHIMADZU, SPM-9700HT). Samples were placed on the AFM sample holder with PBS covering the tissue to maintain hydration. The Young’s modulus (*E*) of the extracellular environment were measured using V-shaped Au-coated silicon nitride cantilever (µmash, XCN12; normal spring constant of *k* = 0.08 N/m) with a spherical polystyrene bead (radius=5 µm; Polysciences, 17136) attached to the tip. The spring constant of each cantilever was calibrated manually prior to use. The effective load to the cantilever (*P*), referred to as “stiffness” in this study, was calculated by fitting the loading force-displacement curve (*F*(*h*)) to the Herz model, using the tip radius (*R*), and indentation depth (*h*), assuming a Poisson’s ratio (*v*) of 0.5 (20). The Herz model formula used in this study is as follows:

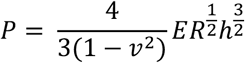

### Decellularization

Decellularization was performed as previously described (40). Briefly, cryosectioned samples were first washed in PBS at room temperature (RT) for 10 minutes to remove residual OCT compound. Samples were then incubated in deionized water for 10 minutes, followed by immersion in 2% (w/v) sodium deoxycholate (SDC) for 1 hour (two consecutive 30-minutes incubations). After SDC treatment, samples were washed in PBS at RT for 5 minutes, and this washing step was repeated three times to ensure complete removal of detergent. All steps were carried out with gentle agitation at 80 rpm using an orbital shaker. Subsequently, samples were incubated in DNase I at 37 °C for 20 minutes, followed by three additional PBS washes (5 minutes each at RT). Decellularized samples were used immediately for either AFM measurement or immunostaining.

### Immunohistochemistry

Immunohistochemistry was performed as previously described (12) using the following primary antibodies: anti-sarcomeric alpha actinin (Abcam, EP2529Y; 1/200) and anti-myosin light chain (Sigma, K36; 1/200).

### Imaging

Immunostained sections were imaged using an Olympus FV3000 confocal microscope (Olympus, Japan). Brightfield images of extracted hearts were acquired using a Leica microscope MZ165FC stereomicroscope equipped with a PLANAPO 1.0x objective and LAS software (version 4.8).

### Indentation

For the indentation experiment, 60 hpf zebrafish embryos were anesthetized with MS-222 (Sigma, A5040) and mounted in 2% low-melting-point agarose (SeqPlaque, 50101) in a supine position. Glass capillaries (Narishige, G-1) were pulled using a needle puller (P-97/IVF, Sutter Instrument) and flame-polished to produce a rounded tip. Prepared glass needles were mounted on a micromanipulator (Narishige, MN-153) and used to apply localized mechanical indentation.

### Statistical Analysis

Statistical analyses for each experiment are described in the corresponding figure legends. All analyses were performed using Prism 9 (GraphPad) and Excel (Microsoft). Data were first assesseds for normality using D’Agostino–Pearson normality test. Since at least one of the datasets being compared was found to be non-normally distributed, non-parametric tests were applied. The Mann–Whitney test was used for comparisons between two groups, and the Kruskal–Wallis test was used for comparisons among more than two groups. n indicates the number of individual data points analyzed, such as AFM measurement or embryos used for *in situ* hybridization.

### Conclusions

In this study, we demonstrated that the BA in teleosts exhibits a uniquely low level of extracellular stiffness compared to the ventricle and the CA in non-teleost fishes.

Loss of *elnb* function or artificial stiffening of the extracellular environment led to increased stiffness and altered cell fate in the BA. These findings suggest that extracellular stiffness plays a regulatory role in cell fate determination within the OFT, thereby contributing to BA development during embryogenesis and to its evolutionary acquisition in teleosts.

## Acknowledgments

We would like to thank Dr. Akinori Kogure (Shimadzu Scientific Instruments, INC.) for helpful discussion about AFM measurement and data fitting. This work was supported by a Grant-In-Aid for Scientific Research (C) (20K06773) to Y.M.

## Author contributions

Y.M. conceived of and conceptualized the project. S.M., Y.I. and R.W. performed the experiments. S.M., R.W. and T.M. analysed the data. Y.M. wrote the manuscript draft. All the authors read and approved the final version of the manuscript.

The authors declare no competing interest.

## SI Appendix Figures

**Fig. S1.**
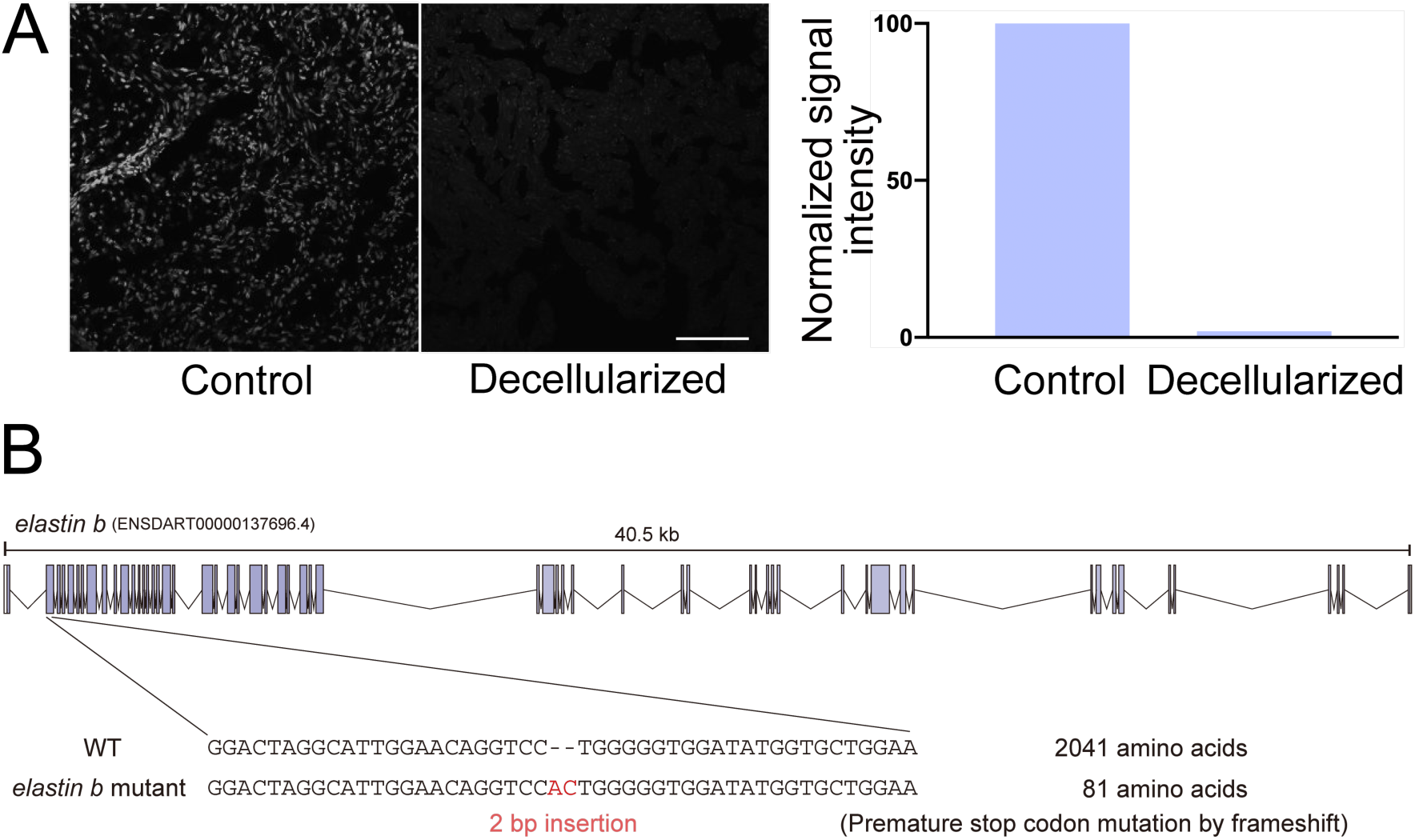
Assessment of decellularization and mutation in *elastin b* knockout mutants. (A) Representative DAPI-stained images of adjacent sections from adult zebrafish heart, comparing a control (non-decellularized) section and a decellularized section. The decellularized sample shows a marked reduction in nuclear signal, indicating effective removal of cellular components. (B) Schematic representation of the CRISPR/Cas9-induced mutation in *elastin b*. A 2-bp insertion in exon 2 introduces a frameshift that results in a premature stop codon, producing a truncated protein of 81 amino acids instead of the full-length Elastin b protein comprising 2041 amino acids.

